# Detection of a biolistic delivery of fluorescent markers and CRISPR/Cas9 to the pollen tube

**DOI:** 10.1101/2021.02.15.431139

**Authors:** Shiori Nagahara, Tetsuya Higashiyama, Yoko Mizuta

## Abstract

In recent years, genome-editing techniques, such as the CRISPR/Cas9 system, have been highlighted as a new approach to plant breeding. *Agrobacterium*-mediated transformation has been widely utilized to generate transgenic plants by introducing plasmid DNA containing CRISPR/Cas9 into plant cells. However, this method is generally applicable to a limited range of plants, such as model species. To overcome this limitation, we developed a method to genetically modify male germ cells without the need for *Agrobacterium*-mediated transfection and tissue culture, by using tobacco as a model. In this study, plasmid DNA containing sequences of Cas9, guide RNA, and fluorescent reporter was introduced into pollen using a biolistic delivery system. Based on the transient expression of fluorescent reporters, the *Arabidopsis UBQ10* promoter was found to be the most suitable for driving expression of the delivered gene in pollen tubes. We also evaluated delivery efficiency in male germ cells in the pollen by expression of the introduced fluorescent marker. Mutations were detected in the target gene in the genomic DNA extracted from CRISPR/Cas9 introduced pollen tubes but were not detected in the negative control. Bombarded pollen germinated pollen tubes on the stigma and produced two sperm cells within the pistil. We also observed ovules showing fluorescence derived from bombarded pollen. The findings of this study provide important insights into the editing of pollen tube genomes and the delivery of genome-modified male germ cells for seed production.

## Introduction

Recently, there has been rapid progress in the development of genome engineering techniques, which have made it possible to perform specific modifications of selected target genes. One of the most widely used methods is targeted genome editing using clustered regularly interspaced short palindromic repeats (CRISPR) and CRISPR-associated protein 9 nuclease (Cas9) (Cong et al. 2013). This method is widely used in various organisms, including plants, because of its suitability for genetic engineering (Li et al. 2013; Nekrasov et al. 2013; Osakabe and Osakabe 2015; Osakabe et al. 2016). In animals, to produce genetically heritable traits of interest, Cas9 protein and sgRNA complexes are delivered into zygotes or eggs, resulting in the highly efficient production of genetically modified animals (Wang et al. 2013). In flowering plants, flowers contain male and female gametophytes, which produce gametes. The female gametophyte of angiosperms is located in the ovary, and the female gamete (egg cell) is deeply embedded within the ovule (Zhou and Dresselhaus 2019). After flowering, pollen grains land on the stigma of the pistil, and they produce cylindrical tip-growing cells, referred to as pollen tubes, which contain two male gametes (sperm cells). Unlike animal cells, angiosperm sperm cells are non-motile, and thus pollen tubes play an essential role in conveying these toward the ovule enclosing an egg cell and a central cell to facilitate double fertilization (Dresselhaus et al. 2016). Pollen tubes serve as vector cells that deliver copies of the male genome to the female gametophyte to produce the seeds that give rise to the next generation. Pollen is easier to access and handle than are egg cells or zygotes, and such features are common to a wide range of angiosperms.

In order to generate genome-modified plants, it is necessary to introduce a CRISPR/Cas9 cassette into plant cells. Methods currently used to deliver material to plant cells can be divided into three broad categories: chemical, physical, and biological (Newell 2000). Among the chemical methods, polyethylene glycol (PEG) can be used to facilitate highly efficient transfection (Toda et al. 2019); however, this approach requires the production of protoplasts, from which it is typically difficult to regenerate plants via callus, and tends to be associated with a high frequency of somaclonal variation (Fossi et al. 2019). Physical delivery methods, such as electroporation, similarly require the production of protoplasts to enable gene transfer (Woo et al. 2015). As a biological technique, *Agrobacterium*-mediated transformation of foreign genes has been the most frequently used in plants (Clough and Bent 1998); however, it has only been applied in a limited range of model plants, for which sufficient infection and culture methods have been established. Similarly, callus induction and/or plant regeneration tend to be difficult in most plant species (Bregitzer et al. 1998). Therefore, it is anticipated that developing a simple and convenient method for introducing CRISPR/Cas9 into a wide range of plant species will contribute to significant advances in plant molecular genetic studies.

Biolistic delivery, also referred to as particle bombardment, can be used to facilitate the transient introduction of exogenous substances into cells (Sanford 2000), and it has been widely used to transfer genes into the cells of a range of plant species (Wang and Jiang 2011). Notably, in recent years, biolistic delivery has been applied to deliver the CRISPR/Cas9 cassette into immature maize embryos (Svitashev et al. 2016; Svitashev et al. 2015) and wheat embryos (Hamada et al. 2017). Given that embryos become mature plants, genetic manipulation of their cells can be effective. However, the embryos thus generated become chimeric plants that necessitate subsequent selection in the resulting progenies. Consequently, there is a need to develop a method that can be used for the simple and efficient delivery of CRISPR/Cas9 cassettes and that is applicable to a range of plant species without the necessity of regeneration steps.

To this end, in the present study, we developed a method for the biolistic delivery of CRISPR/Cas9-harboring plasmid DNAs into plant pollen for engineering the genome of male gametophytes. Pollen grains are simple structures containing male germ cells, which are suitable for the introduction of exogenous material, as has been demonstrated recently with various approaches (Bhowmik et al. 2018; Eapen 2011; Zhao et al. 2017). We identified a suitable promoter for the control of gene expression in the pollen of four angiosperms following particle bombardment and assessed the efficiency with which genes were introduced into pollen using this technique. As a consequence of introducing plasmid DNA containing CRISPR/Cas9, we detected certain mutations within the target sequence in genomic DNA extracted from both leaves and pollen tubes. Additionally, we observed that bombarded pollen germinated and delivered generative or sperm cells and reached the ovules *in vivo*. These results will contribute to the production of genome-modified plants based on pollination with genome-edited pollen obtained using a biolistic delivery system.

## Materials and Methods

### Plant materials

Seeds of *Nicotiana tabacum* cv. ‘SR1’ were obtained from the Leaf Tobacco Research Center. *Nicotiana benthamiana*, *N. tabacum* ‘SR1’, and *Torenia fournieri* cv. ‘Blue and White’ plants were grown in soil in a green room at 25–30°C under long-day conditions (16 h light/8 h dark), and *Solanum lycopersicum* cv. ‘Micro-Tom’ plants were grown in soil in a greenhouse at 20–30°C. Young leaves and mature pollen from newly opened flowers were used for bombardment.

### Plasmid construction

The plasmid vectors used for particle bombardment in this study are listed in Table S1. The constructs *LAT52p::mApple* (YMv32), *LAT52p::Venus* (YMv35), *LAT52p::mTFP1* (YMv37), and *AtRPS5Ap::H2B-tdTomato* (DKv277) were obtained from previous studies (Adachi et al. 2011; Mizuta et al. 2015). *35Sp::H2B-mClover* (DKv700) and *35Sp::H2B-tdTomato* (DKv744) were provided by Dr. Daisuke Kurihara, and *35Sp::mTFP1* (DKv327) was provided by Dr. Noriko Inada. The *AtRPS5Ap::sGFP* vector (sSNv10), in which the *sGFP* gene is driven by the *Arabidopsis thaliana RPS5A* (*RIBOSOMAL PROTEIN SUBUNIT 5A; At3g11940*) promoter, was produced by inverse PCR of the *AtRPS5Ap::H2B-sGFP* vector (Maruyama et al. 2013) and self-ligation of the PCR product to remove the *H2B* (*HISTONE 2 B; At1g07790*) sequence. The sequences of *AtUBQ10p::sGFP* (sSNv25), *AtUBQ10p::tdTomato* (sSNv26), and *AtUBQ10p::H2B-mClover* (sSNv28) followed by the *Nos*-terminator sequence were isolated from the DKv909, DKv922, and DKv916 vectors, respectively (Kurihara et al. 2015), by *Hin*dIII/*Eco*RI digestion. Each fragment was cloned into the pGreen0029 vector (Hellens et al. 2000) using *Hin*dIII and *Eco*RI sites. The CRISPR/Cas9 vector targeting the *NbPDS3* gene (sSNv21) was constructed based on a previous study (Tsutsui and Higashiyama 2017). The *AtRPS5A* promoter was replaced with the *AtUBQ10* (*UBIQUITIN 10; At4g05320*) promoter to drive Cas9 gene expression, and the target sequence for the *NbPDS3* gene was then introduced via *Aar*I digestion (sSNv21). The primers used for plasmid construction are listed in Table S2.

### Plant transformation

The sSNv28 vectors were introduced into *Agrobacterium tumefaciens* strain LBA4404 harboring the pSoup plasmid (Hellens et al. 2000) by electroporation, and *N. benthamiana* leaf discs were infected with *Agrobacterium*. The infected leaf discs were cultured on callus induction medium (1× Murashige and Skoog Basal Medium, 3% (w/v) sucrose, 0.8% (w/v) Bacto agar, adjusted to pH 5.8 with KOH] containing 0.05 mg/L 1-naphthaleneacetic acid, 0.5 mg/L 6-benzylaminopurine, 100 mg/L kanamycin sulfate, and 300 mg/L cefotaxime sodium. The plants that regenerated from calli were transferred to medium lacking hormones to induce root germination and were eventually transferred to soil.

### Calculating the percentage area of pollen nuclei to cytoplasm

To calculate the percentage of area of pollen nuclei to the cytoplasm, we analyzed fixed raw and raw pollen grains. Wild-type pollen grains were fixed with a 9:1 mixture of ethanol and acetic acid (v/v) for 10 min, and the samples were directly stained with DAPI solution and incubated for more than 10 min. After staining, the pollen grains were washed twice with water, mounted on glass slides, and observed under an inverted fluorescence microscope (Eclipse Ti2; Nikon, Tokyo, Japan). The pollen from *UBQ10p::H2B-mClover* was collected in a liquid pollen germination medium [0.01% (w/v) boric acid, 1 mM CaCl_2_, 1 mM Ca(NO_3_)_2_, 1 mM MgSO_4_, and 10% (w/v) sucrose] adjusted to pH 6.5 with KOH (Wang and Jiang 2011), and the sample was immediately observed under a fluorescence microscope. ImageJ software (https://imagej.nih.gov/ij/index.html) was used to calculate generative nucleus to cytoplasm (GN/C) ratio of 20 pollen grains.

### Biolistic delivery of plasmid DNA

The gold particles (0.6-μm diameter) used for biolistic delivery were washed with absolute ethanol, rinsed twice with sterilized water, and suspended in sterilized water to prepare a 30 mg/mL gold solution. The gold solution was dispensed (10 μL per shot) and mixed with 200–1,000 ng of plasmid DNA(s) per shot in an agitating mixer, to which 4 μL 0.1 M spermidine and 10 μL 2.5 M calcium chloride per 10 μL of the gold solution were subsequently added. The resulting DNA-coated gold particles were collected by centrifugation at 3,300 *g* for 30 s. The DNA-coated gold particles were then washed once with 70% ethanol, twice with absolute ethanol, and resuspended in absolute ethanol (10 μL per shot). Particle bombardment was performed using a PDS-1000/He system (Bio-Rad Laboratories, USA). The distance between the macro-carrier and target cells was adjusted to approximately 3.0 cm, the helium gas pressure was set to 1,100 psi, and the degree of vacuum was set to at least −25 inHg. For leaf bombardment, we used the leaves of mature *N. benthamiana* plants, which were taped to a plastic petri dish. The cut ends of the bombarded leaves were covered with a wet wipe and cultured at 25–30°C for 20– 24 h under humid, dark conditions. For pollen bombardment, pollen was collected immediately prior to bombardment and distributed on pollen germination medium solidified with 1% (w/v) NuSieve GTG Agarose (Lonza, Switzerland). Pollen germination media for *Nicotiana* (Wang and Jiang 2011) and torenia (Okuda et al. 2009) were used as described previously. The same pollen germination medium as that used for *Nicotiana* was used. After being bombarded, the treated pollen was cultured directly on the medium and observed under an inverted fluorescence microscope (Eclipse Ti2, Nikon; AXIO imager A2, Zeiss). Biolistic delivery into leaf and pollen was examined twice, and gene introduction was evaluated by fluorescence expression.

### Detection of Cas9-induced genome editing

In addition to the CRISPR/Cas9 vector described above, plasmid vectors encoding fluorescence protein markers were simultaneously coated with gold particles. Bombarded leaves were observed under a fluorescence microscope (Eclipse Ti2, Nikon; AXIO imager A2, Zeiss) to confirm the efficiency of delivery. An area of approximately 1.5 cm in diameter around the blast center was collected, immersed in a DNA extraction buffer [0.2 M Tris-HCl (pH 8.0), 0.25 M NaCl, 25 mM EDTA, and 0.5% (w/v) SDS], and frozen in liquid nitrogen prior to subsequent analysis. Genomic DNA was extracted from the leaf material using a homogenizer and sonicator. Pollen samples with or without fluorescent signals were collected together into the same DNA extraction buffer and the genomic DNA was then extracted. For PCR analysis, the following steps were performed according to Nekrasov et al. (2013). Genomic DNA was digested with *Mly*I and PCR-amplified with the primer pair *PDS*_*Mly*IF and *PDS*_*Mly*IR (Nekrasov et al. 2013; Table S2). The PCR product thus obtained was subsequently digested with *Hin*fI and/or used directly as a nested PCR template to remove non-specific DNA fragments. The nested PCR product was cloned into the pCR-BluntII-TOPO vector (Thermo Fisher Scientific, USA) and introduced into *Escherichia coli* Mach1 T1^R^ competent cells (Thermo Fisher Scientific). Colonies possessing the fragment of the mutated *NbPDS3* gene were identified by colony PCR using primers M13 forward, M13 reverse, and *NbPDS3*_primer-m (Table S2), of which the 3ʹ end was fully matched to the wild-type *NbPDS3* sequence. Accordingly, for colonies containing mutated *NbPDS3* fragments, few or no bands were amplified using *NbPDS3*_primer-m. A mutation in the *NbPDS3* gene was confirmed by sequence analysis of the plasmid vectors extracted from individual colonies.

### Aniline blue staining of pollinated pistils

Wild-type *N. benthamiana* and *N. tabacum* pistils were emasculated 2 days prior to pollination. The pollen spread on germination medium that had been hydrated for 15 min was used to pollinate the emasculated pistils using a dissecting needle. At 24 h after pollination, the pollinated flowers were collected, and the remaining petals and sepals were removed. The pistil was fixed with a 3:1 mixture of ethanol and acetic acid (v/v) overnight and treated with 1 N NaOH solution for 1 day. Thereafter, the pistils were stained with 0.1% (w/v) aniline blue in 0.1 M K_3_PO_4_ buffer for 1 day. Images were obtained using an Eclipse Ti2 fluorescence microscope under UV light.

### Semi-*in vivo* pollen tube growth assay using *N. tabacum*

*N. tabacum* pistils were emasculated 2 days prior to pollination. Plasmid vectors encoding *UBQ10p::sGFP* and *35Sp::H2B-tdTomato* were simultaneously coated on gold particles (mixture 1), as were the vectors encoding *LAT52p::mApple* and *UBQ10p::H2B-mClover* (mixture 2). Immediately prior to spreading onto the macro-carrier, mixtures 1 and 2 were mixed and then introduced into pollen via a single bombardment shot. Twenty-four hours after pollination, the pollinated style was excised with a razor 5 mm above the ovary and placed horizontally on the aforementioned pollen germination medium solidified with 1% (w/v) NuSieve GTG Agarose (Lonza), followed by incubation at 25–30°C for 20 h under humid, dark conditions. Pollen tubes emerging from the cut end of the pistil were observed using an Eclipse Ti2 fluorescence microscope.

### *In vivo* experiments using *N. tabacum*

*N. tabacum* pistils were emasculated 2 days prior to pollination. Pollen was bombarded with the aforementioned mixture 1 and used immediately thereafter to pollinate the emasculated pistils. At 24 h post-pollination, the pollinated style was cut longitudinally using a razor, and at 24 h and 48 h post-pollination, the ovary wall was removed with forceps to facilitate observation of the pollen tube and ovule within the ovary. The samples were placed in 10% glycerol (v/v) and observed under an Eclipse Ti2 fluorescence microscope.

## Results

### The *AtUBQ10* promoter is suitable for driving transient gene expression in pollen germ cells

In order to identify an effective promoter for driving the expression of Cas9 in pollen, we evaluated the activity of selected promoters in bombarded pollen based on fluorescent protein expression. Fluorescent proteins were driven under the control of cauliflower mosaic virus (CaMV) 35S*, A. thaliana* Ribosomal protein S5A (*AtRPS5A*), and *A. thaliana* UBIQUITIN10 (*AtUBQ10*) promoters, which have previously been used as constitutive promoters (Liang et al. 2018; Nekrasov et al. 2013; Tsutsui and Higashiyama 2017), as well as the pollen vegetative cell-specific *LAT52* promoter derived from *S. lycopersicum* (Eady et al. 1995). The plasmid DNA vectors containing these promoters, which were introduced into pollen and leaves, are summarized in Table S1. Tri-cellular pollen is generally short-lived relative to bicellular pollen, which increases the difficulty of handling this type of pollen (Hoekstra and Bruinsma 1975). In the present study, we used four species with bi-cellar pollen to investigate the introduction of the aforementioned plasmid DNA vectors, namely, *N. benthamiana* (tobacco), *N. tabacum* (tobacco), *T. fournieri* (torenia), and *S. lycopersicum* (tomato), for which the *in vitro* induction of pollen tube germination has been established (Liang et al. 2018; Okuda et al. 2009; Paungfoo-Lonhienne et al. 2010; Zhang et al. 2020). Biolistic delivery into leaf and pollen was examined twice, and gene introduction was evaluated by fluorescence expression. Accordingly, we observed that in the bombarded leaves of all examined species, fluorescent proteins were expressed under the control of each of the three constitutive promoters (Fig. 1), whereas no fluorescent proteins were detected in leaves containing the pollen-specific *LAT52* promoter (Table 1). In the case of bombarded pollen, *UBQ10* and *RPS5A* promoters were found to drive H2B-fused fluorescent protein expression in all examined species (Fig. 1), whereas the *AtRPS5A* promoter showed low activity, with fluorescence being detected only in the pollen cell nuclei of *N. benthamiana* and *N. tabacum* (Table 1, Fig. 1). Similarly, the 35S promoter was active in the pollen of *N. benthamiana* and *N. tabacum*, but not in that of torenia or tomato (Table 1). A promoter that controls stable gene expression in generative cells is most suitable, as these cells contain the genome that is transmitted to the next generation. Accordingly, we used the *AtUBQ10* promoter to drive Cas9 in pollen transformed via particle bombardment. Furthermore, we used tobacco pollen in subsequent bombardment assays, as large amounts of pollen can be obtained from these plants.

**Fig. 1.**
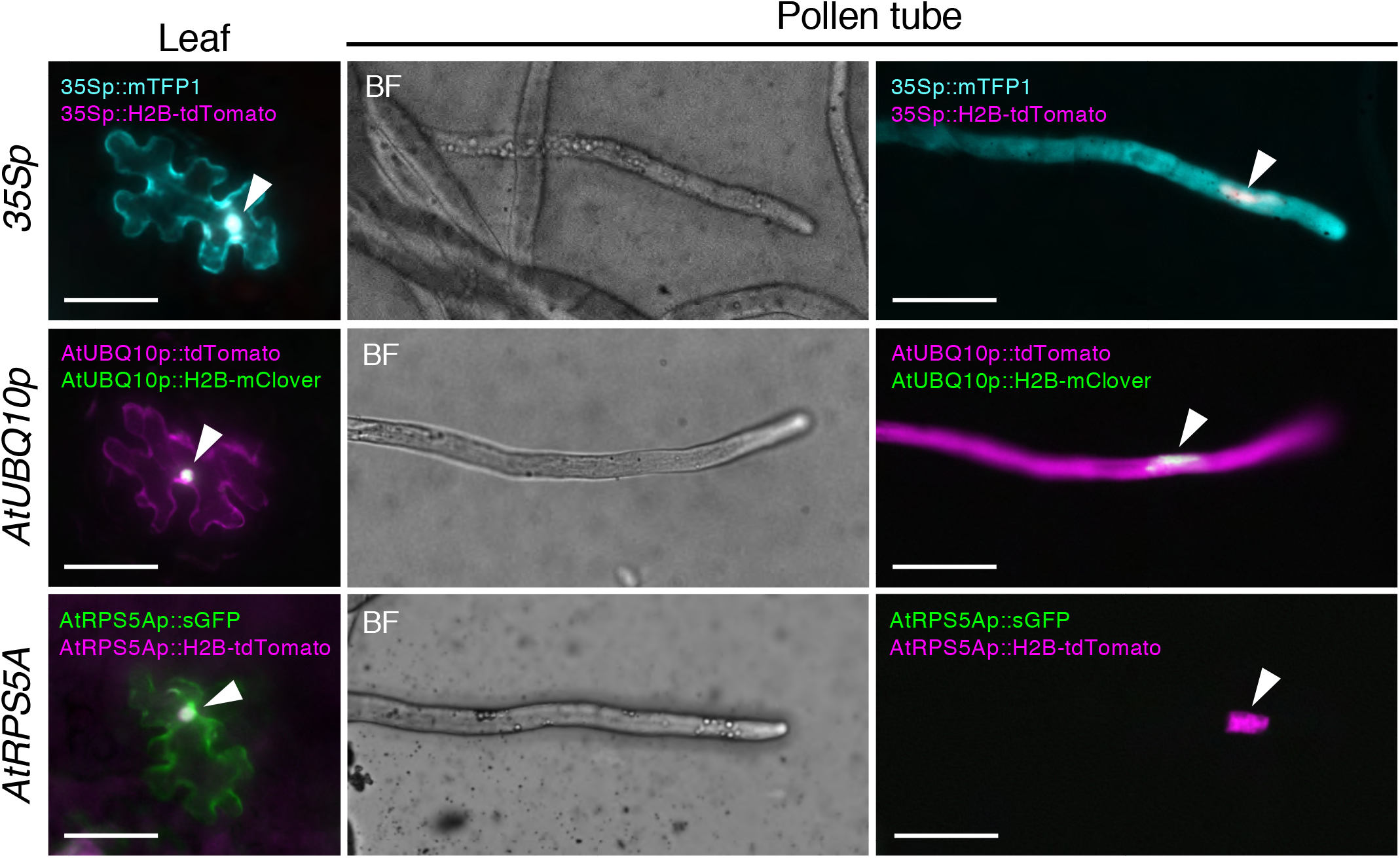
Promoter activity in *Nicotiana benthamiana*. CaMV 35S (35S), *Arabidopsis thaliana (At) UBQ10* and *AtRPS5A* promoter activity in *N. benthamiana* leaves and pollen tubes. The epidermal cells in *N. benthamiana* leaves show the cytosolic and nuclear expression of fluorescent proteins driven by the 35S, *AtUBQ10*, or *AtRPS5A* promoter. Note that, as fluorescent proteins showing cytosolic expression were also localized in nuclei, pseudo-colors appear to be white in the nuclei (arrowheads). The *N. benthamiana* pollen tubes show the signals of fluorescent proteins driven by *35S* and *AtUBQ10* promoters in both cytosol and nucleus (arrowheads), whereas the signals of fluorescent proteins driven by the *AtRPS5A* promoter were observed only in the pollen tube nuclei in the merged fluorescent images. The plasmid DNAs used in the experiment are listed in Table S1. BF, bright field. Scale bars: 50 μm.

**Table 1.**
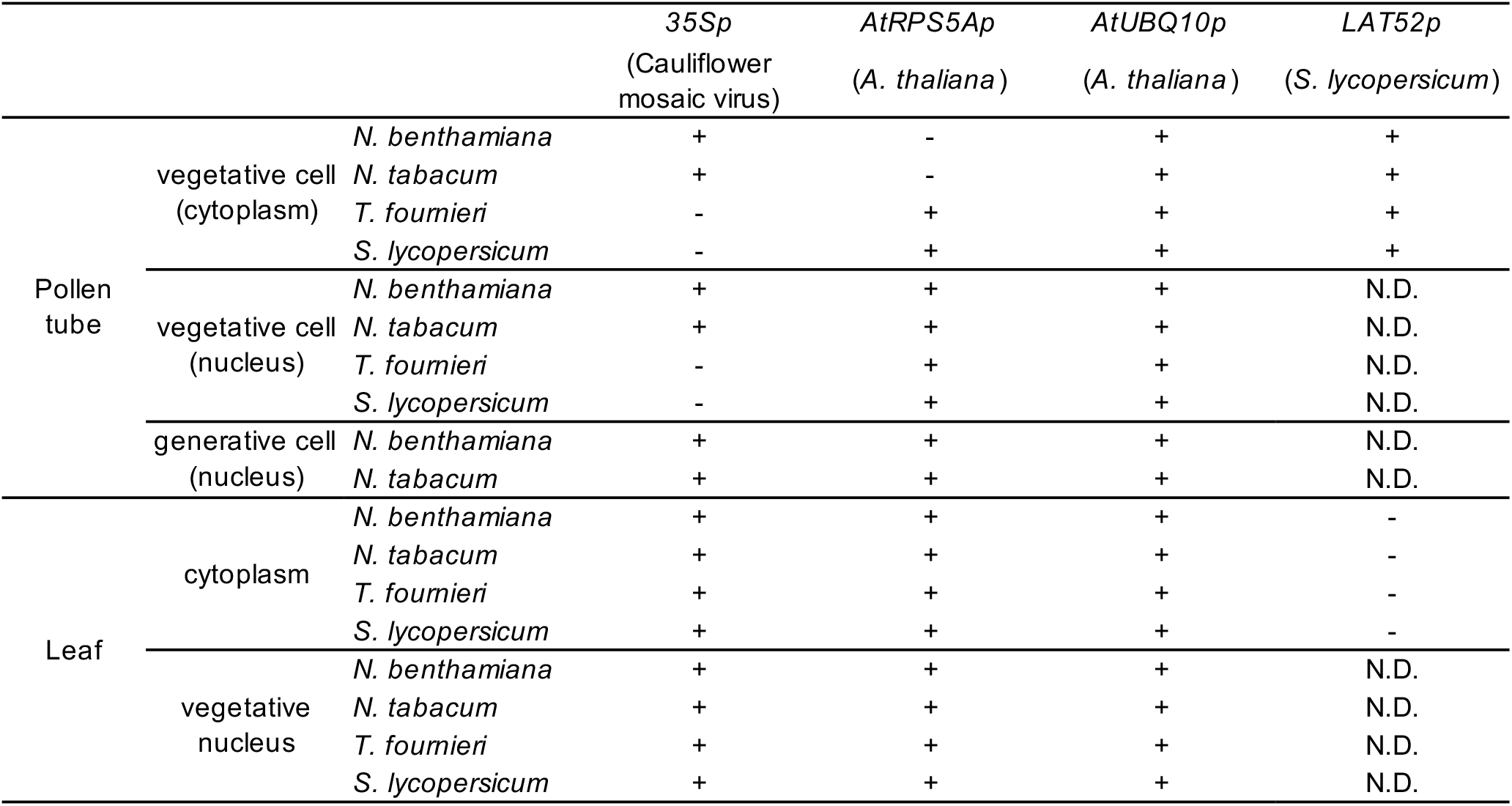
Transient expression of the fluorescent proteins by the biolistic delivery of plasmid DNA

### Efficiency of gene delivery into generative cells by particle bombardment

In general, the efficiency with which genes are delivered by particle bombardment is typically as low as several percent (Wang and Jiang 2011). Given that genome editing occurs in only a fraction of the gene-bombarded pollen, we initially investigated the efficiency with which genes are delivered into pollen. Plasmid DNA vectors containing fluorescent protein driven under the control of the aforementioned promoters were introduced into the pollen of *N. benthamiana* by particle bombardment. We simultaneously introduced two types of plasmid DNA encoding mApple and H2B-mClover, respectively, into the same pollen, and, accordingly, observed mApple signals in the cytoplasm, whereas mClover signals were localized in the nuclei, indicating that subcellular structures in pollen and pollen tubes were labeled transiently (Fig. 2a). Moreover, the observed expression of delivered genes in the vegetative and generative cells of the germinated pollen tubes indicated that the gold particles had penetrated into the generative cells enclosed within the pollen (Fig. 2b). Interestingly, in some pollen tubes, we observed dot-like structures, which are assumed 0.6 μm gold particles (Fig. 2c). Furthermore, following bombardment, some pollen tubes were found to contain two sperm cells that were divided from the generative cell (Fig. 2d). Although nuclear signals of both vegetative and sperm cells were observed, fluorescence signals derived from the delivered plasmid DNA were detected only in the vegetative cells, indicating that gold particles were introduced only into the vegetative cells of pollen, and that generative cell division occurred to produce two sperm cells.

**Fig. 2.**
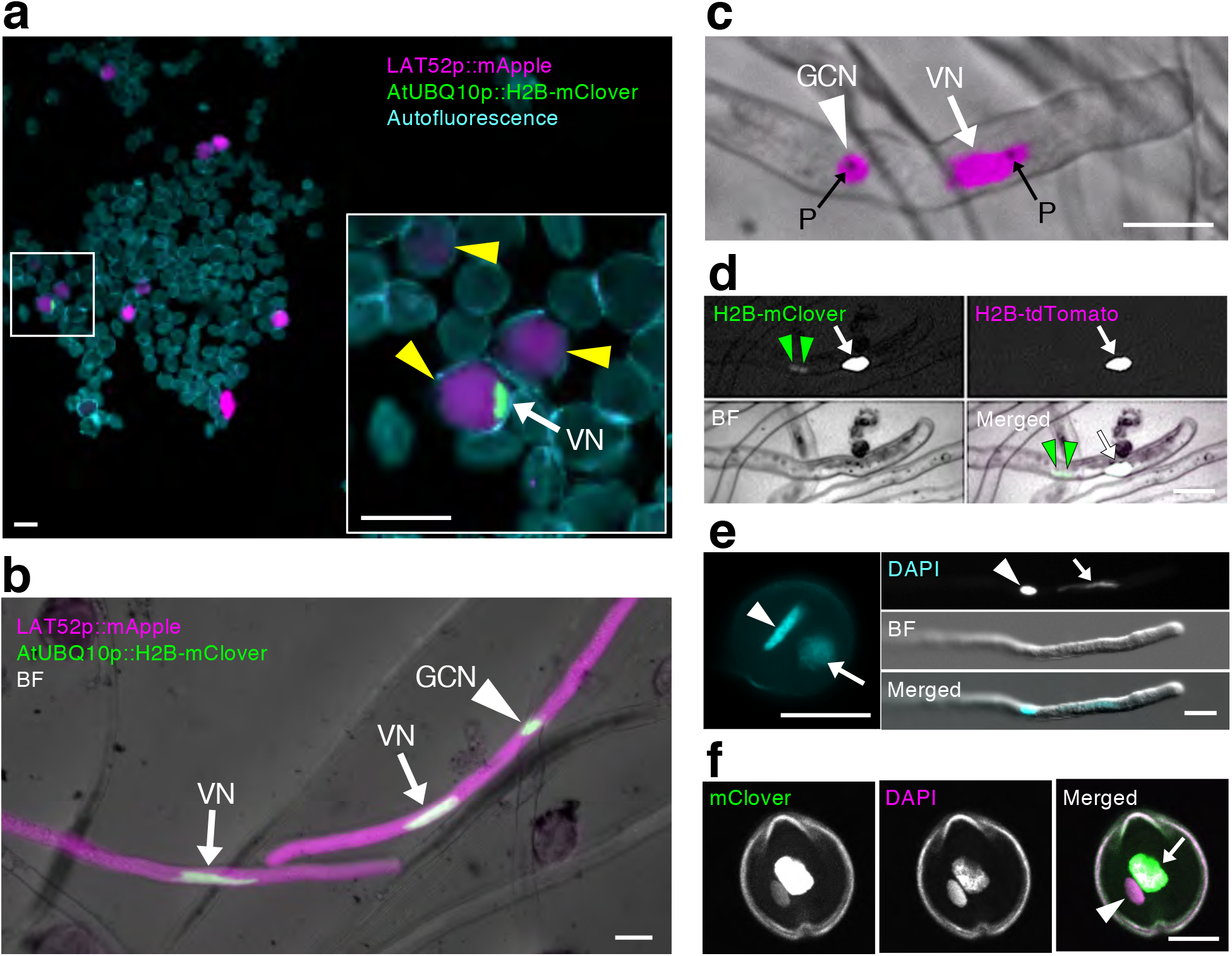
Biolistic delivery of plasmid DNAs into the vegetative and generative cells of *Nicotiana benthamiana* pollen and pollen tubes. **a:** Bombarded pollen expressing fluorescent proteins at 6 h after bombardment. The inset shows a magnified view of the area demarcated by the white-bordered square. Yellow arrowheads indicate mApple-positive pollen cytosol (magenta; *LAT52p::mApple*) and the white arrow indicates a mClover-positive vegetative cell nucleus (green; *AtUBQ10p::H2B-mClover*). The outline of pollen grains was visualized by auto-fluorescence (cyan). **b:** Pollen tubes germinated from bombarded pollen at 20 h after bombardment. mApple and mClover fluorescence and bright-field (BF) images are merged. The arrows and arrowhead indicate mClover signals in vegetative nuclei and a generative cell nucleus, respectively. The right-hand pollen tube shows mClover signals in both vegetative and generative cell nuclei. **c:** Pollen tubes germinated from pollen bombarded with *35Sp::H2B-tdTomato* plasmid DNA. RFP fluorescence and BF images are merged. An arrow and an arrowhead indicate H2B-tdTomato signals in a vegetative nucleus and a generative cell nucleus, respectively. Delivered dot-like gold particles are indicated by black arrows. **d:** Germinated pollen tubes of *AtUBQ10p::H2B-mClover*, containing two sperm cells at 10 h after bombardment with *35Sp::H2B-tdTomato* plasmid DNA. Green arrowheads indicate two sperm cell nuclei and arrows indicate a vegetative nucleus. A green signal was detected in the nuclei of both vegetative and sperm cells, whereas a magenta signal was observed only in the vegetative cell. **e:** A DAPI-stained fixed pollen grain and pollen tube of *N. benthamiana*. The arrow and arrowhead indicate the vegetative and generative cell nucleus, respectively. **f:** DAPI-stained unfixed pollen derived from *AtUBQ10p::H2B-mClover* transgenic *N. benthamiana*. The arrow and arrowhead indicate the vegetative and generative cell nucleus, respectively. VN, vegetative nucleus; GCN, generative cell nucleus; P, gold particles within the generative cell and pollen tube. Scale bars: **a:** 50 μm, **b−e:** 20 μm, **f**:10 μm.

To obtain an estimate of the efficiency with which genes are delivered into the vegetative and generative cells of pollen, we subsequently determined the ratio of transformed pollen grains to total pollen grains. The estimated percentages of the vegetative cells of transformed pollen showing expression of mApple in the cytoplasm and mClover in vegetative nuclei were 1.2% (18.3 ± 6.5 pollen grains) and 1.1% (16.0 ± 4.7 pollen grains), respectively (1,479 ± 298.5 pollen grains per experiment, n = 3). This indicated that regardless of protein localization (i.e., cytoplasm or nucleus), there is little difference in the efficiency of gene delivery and expression. However, we estimated that only 0.2% of the transformed pollen showed mClover signals in generative cell nuclei, and in such pollen, both vegetative and generative nuclei were labeled (3.5 ± 1.0 pollen grains). This indicates that approximately one-sixth of the delivered pollen was also introduced into the generative cells. Staining of the nuclei of wild-type pollen indicated that the ratio of pollen generative nucleus to cytoplasm (GCN/C) was 4.8% ± 0.5% (n = 20 pollen grains) (Fig. 2e). We also produced *Agrobacterium*-mediated transgenic *N. benthamiana* expressing *H2B-mClover* under the control of the *AtUBQ10* promoter and detected the corresponding H2B-mClover signals in the nuclei of both vegetative and generative cells (Fig. 2f). Similar results were also obtained using nuclear-labeled marker lines transformed with *UBQ10p::H2B-mClover* without fixation (3.9% ± 1.0%, n = 14 pollen grains), thereby indicating that more genes had been introduced into the generative cells than expected based on the GCN/C area ratio of pollen. Based on these observations, we concluded that particle bombardment is applicable for gene delivery into vegetative and generative cells, although the efficiency of delivery into the latter was approximately six times lower than that into the former.

### Genome editing in pollen can be induced by exogenous CRISPR/Cas9 components

To facilitate CRISPR-Cas9-mediated genome editing, the introduced gene must initially be transcribed and translated, which results in a time lag before the effects can be detected. Therefore, we performed time-lapse imaging of the bombarded pollen to estimate the length of time required prior to detection of the transient expression of introduced DNA *in vitro*. We found that fluorescent signals of mClover encoded by the *AtUBQ10p::H2B-mClover* plasmid (sSNv28; Table S1) appeared at 2.5–3.5 h after bombardment, whereas those of tdTomato encoded by the *AtUBQ10p::tdTomato* plasmid (sSNv26; Table S1) appeared at 3.5–4.5 h (Fig. 3a, Video S1). In this regard, it has been established that pollen tube germination in *N. benthamiana* commences after 0.5–5 h on the agarose media, and thereafter grows for approximately 24 h (Paungfoo-Lonhienne et al. 2010). Accordingly, we decided to analyze CRISPR/Cas9 plasmid-mediated genome editing in *N. benthamiana* pollen at 20–24 h post-bombardment. To investigate whether the biolistically delivered CRISPR/Cas9 system is functional in pollen, we introduced a plasmid DNA consisting of a human codon-optimized Cas9 gene and an *A. thaliana U6.26* promoter (*AtU6.26*)-driven sgRNA cassette (Tsutsui and Higashiyama 2017). Functioning of the *AtU6.26* promoter and sgRNA in *N. benthamiana* leaves were confirmed by transient expression via agro-infiltration (Nekrasov et al. 2013). We initially introduced *AtUBQ10p::Cas9/U6.26p::NbPDS3-sgRNA* plasmid (sSNv21; Table S1) into *N. benthamiana* leaves by particle bombardment, and then identified different mutation patterns based on sequence analysis (Fig. 3b−3d). Analysis of the sequences obtained from 116 clones derived from the PCR product shown in lane 2 of the gel depicted in Fig. 3b revealed the presence of indels in 17 of these clones. These mutations could be grouped into five different types: 9, 4, and 2 bp deletions or 1 bp insertions of either T or A (Fig. 3d). We subsequently delivered the same plasmid into *N. benthamiana* pollen. Bulk pollen tubes with and without fluorescent signals (Fig. 3c) were collected together, and genomic DNA derived from both vegetative and generative cells of the pollen tubes was extracted. Sequence analysis of 168 clones derived from the PCR product revealed the presence of deletions or substitutions in 33 of these clones (Fig. 3d), which could be grouped into three different types: 1-bp deletion or substitutions of either C to T or T to C. No mutations were detected in genomic DNA extracted from pollen tubes bombarded without plasmid DNA. These results indicate that transient expression of plasmid DNA, including CRISPR/Cas9, delivered via particle bombardment, can induce genome editing in the leaves and pollen of *N. benthamiana*.

**Fig. 3.**
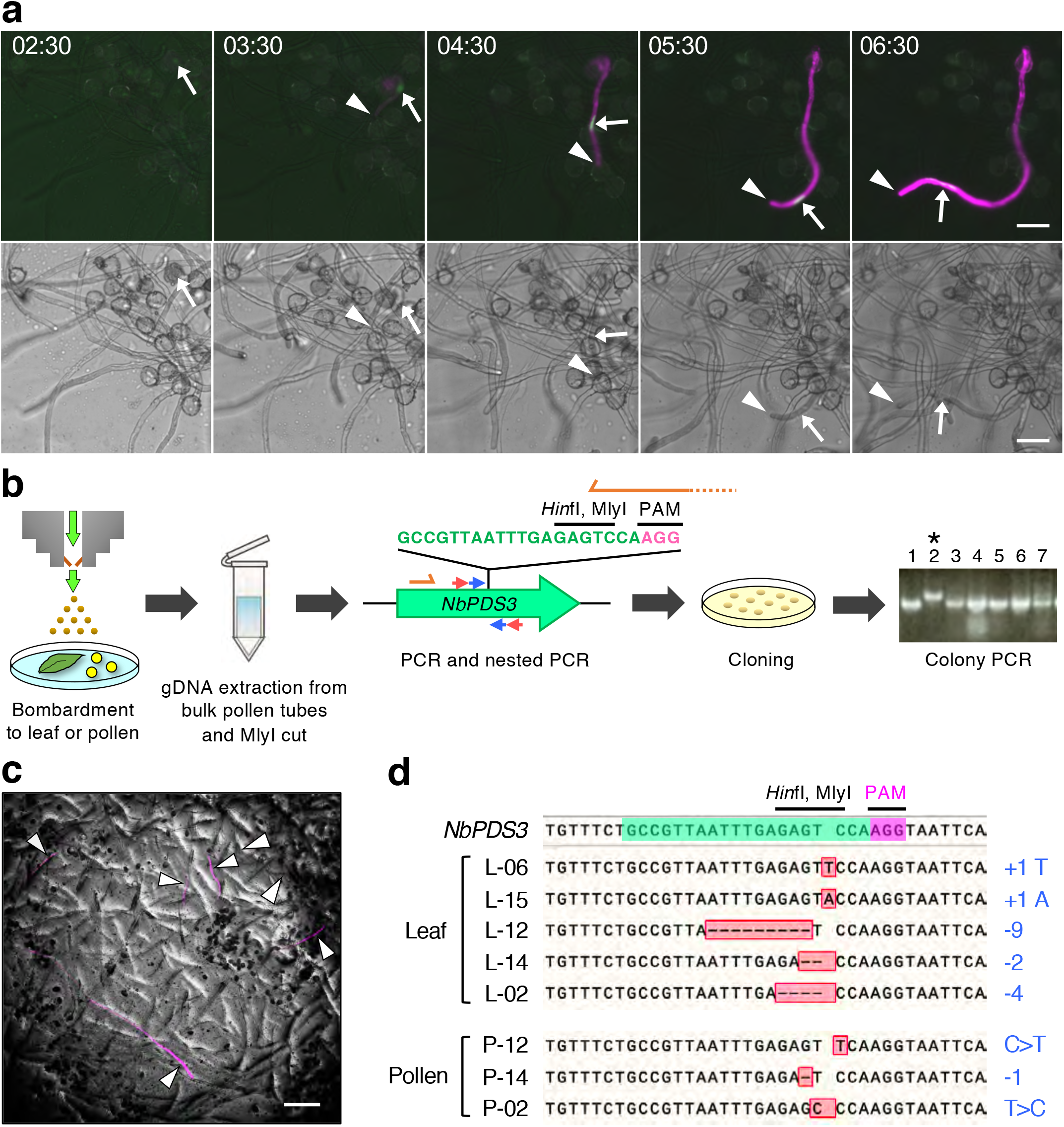
Cas9-induced genome editing via biolistic delivery of CRISPR/Cas9 plasmid DNA introduced into *Nicotiana benthamiana* leaves and pollen. **a:** Time-lapse images of fluorescent protein expression in bombarded pollen. Fluorescent (upper panels) and bright field (BF) (lower panels) images are shown. The signals of tdTomato and H2B-mClover proteins driven by the *AtUBQ10* promoter are shown as magenta and green, respectively. The times denote the period elapsed post-bombardment (hours:minutes). The arrows and arrowheads indicate mClover in the nucleus and the tip of pollen tube of the bombarded pollen. See also Viedeo S1. **b:** Schematic representation of the experimental procedure used to identify mutations in the leaves or pollen into which CRISPR/Cas9 plasmid DNA has been introduced. The bombarded leaves or bulk pollen tubes were transferred to a microtube containing DNA extraction buffer, and genomic DNA was then extracted. The genomic DNA was digested with *Mly*I, and a part of the *N. benthamiana PDS3* (*NbPDS3*) gene was PCR amplified using a primer pair represented by red arrows. Nested PCR was performed directly or after *Hin*fI digestion using a primer pair represented by blue arrows. The partial sequence of the *NbPDS3* gene contains an sgRNA-targeting sequence (green). The PAM sequence is shown in magenta. The nested PCR products were cloned into a cloning vector and transformed into *Escherichia coli*. Colony PCR was performed using primer sequences from the *NbPDS3* gene shown as orange arrows. In the gel image showing PCR bands, only the product shown in lane 2 is mutated. **c:** Pollen tubes derived from bombarded pollen expressing tdTomato at 18 h after bombardment (arrowheads). Genomic DNA was extracted from bulk pollen tubes. **d:** Sequence analysis of the mutation in bombarded leaves and pollen. The upper sequence is the wild-type *NbPDS3* gene. The sequences amplified from both bombarded leaf and pollen containing the CRISPR/Cas9 vector showed mutations in the *NbPDS3* gene. The changes in length and sequence are shown to the right. Scale bars: **a:** 50 μm, **c:** 200 μm.

### Visualization of fertilization process of bombarded pollen *in vivo*

To investigate whether bombarded pollen retains fertilization capacity *in vivo*, we used a nuclear-labeled marker line to investigate pollen tube germination and the delivery of generative cells. For bombardment, we used pollen derived from a *UBQ10p::H2B-mClover* transformant into which plasmid DNA containing the *UBQ10p:H2B-tdTomato* sequence (DKv277; Table S1) was introduced. Thereafter, we performed time-lapse imaging of pollen tube elongation for 18 h after bombardment. Time-lapse observation revealed the delivery of bombarded generative cells within the elongating pollen tube (Fig. 4a, Video S2). It is known that in some species, pollen is unable to germinate on the stigma when the pollen has initially been hydrated on the medium (Zuberi and Dickinson 1985). Thus, we investigated whether *N. benthamiana* and *N. tabacum* pollen that had been hydrated on germination medium could germinate on the pistil. Following hydration, the pollen was immediately used to pollinate emasculated stigmas within 15 min. The pollinated pistils were then fixed and stained with aniline blue solution, and observations indicated that the hydrated pollen of both *N. benthamiana* and *N. tabacum* can germinate on the respective pistils *in vivo* (Fig. 4b, 4c). To enable direct observations of bombarded pollen tube elongation, we conducted a semi-*in vivo* pollen tube growth assay (Palanivelu and Preuss 2006). Owing to the thin style, the pistils of *N. benthamiana* are difficult to dissect without causing damage; therefore, we used *N. tabacum*, which has thick hard pistils, for this assay. Wild-type pollen bombarded with a mixture of plasmid DNAs encoding fluorescent proteins driven by *AtUBQ10* or 35S promoters were immediately used to pollinate emasculated pistils, and pollen tubes were observed, including those derived from bombarded pollen, emerging from the end of the cut style (Fig. 4d). Moreover, we found that the vegetative nucleus in the pollen tube was labeled, and that the lengths of the emerging pollen tubes derived from bombarded pollen were similar to those of other pollen tubes that had germinated from non-bombarded pollen. Furthermore, the fertilization process of bombarded pollen after pollen tube germination was observed *in vivo* (Fig. 5). Emasculated wild-type *N. tabacum* pistil was pollinated with bombarded pollen with plasmid DNA, including *AtUBQ10p::sGFP* and *35Sp::H2B-tdTomato* sequences. Dissection of pistils at 24 h after pollination (Fig. 5a) revealed that the pollen tubes derived from bombarded pollen had elongated within the longitudinal section of the style *in vivo* (Fig. 5b). Moreover, we observed that some pollen tubes had undergone cell division, resulting in the production of two sperm cells and a single vegetative cell within the pollen tube (black arrows in Fig. 5b). Additionally, the pistil was also dissected 48 h after pollination. We observed that bombarded pollen elongated on the placenta in the ovary (Fig. 5c, 5d), and that some ovules had received pollen tubes derived from bombarded pollen grains at 48 h after pollination (Fig. 5e). When bombardment was performed twice and pollinated two flowers each, one, two, three, and five ovules showed fluorescence signals in each ovary at 48 h after pollination. Fluorescence signals derived from pollen tube cytoplasm reporters were detected as a large spot of fluorescence signal inside ovules (Fig. 5f). This suggested that the pollen tubes released their contents, which were detected as a conveniently marking targeted ovules (Palanivelu and Preuss 2006). These observations indicate that even *in vivo* bombarded pollen can germinate and give rise to pollen tubes that undergo subsequent elongation and can deliver their contents, including sperm cells, into the ovule.

**Fig. 4.**
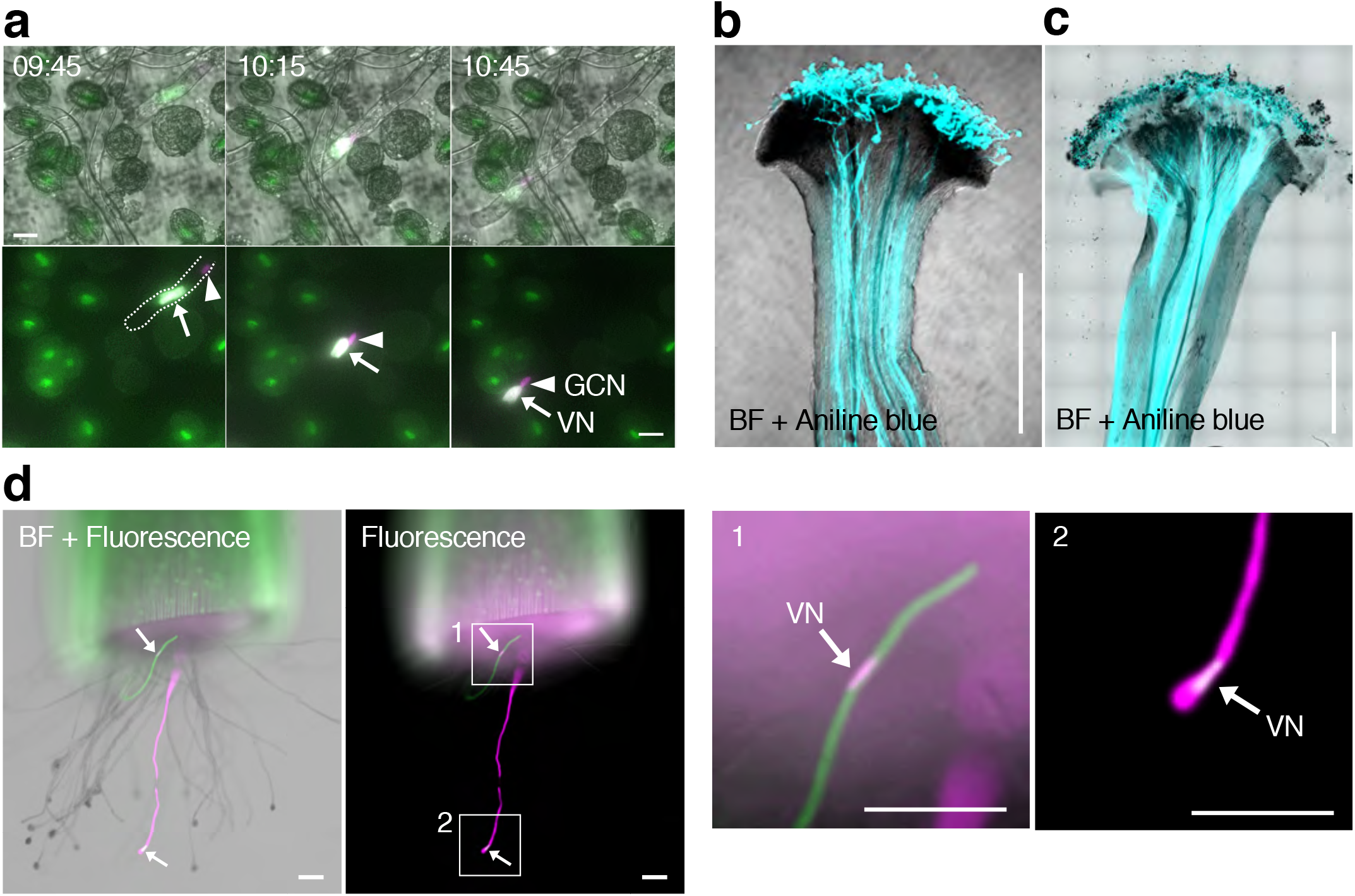
Pollen tubes derived from pollen bombarded *in vitro* and semi-*in vivo* experiments. **a:** Delivery of the bombarded generative cell in a pollen tube of a transgenic *Nicotiana benthamiana* plant. The vector *35Sp::H2B-tdTomato* was introduced into pollen from *AtUBQ10p::H2B-mClover*. Note that the generative cell nucleus appears magenta in color, whereas the vegetative nucleus appears white, due to the lower level of the background expression of *H2B-mClover* in the GCN than VN of the transgenic line, as shown in Fig. 2d, 2f. Upper panels are merged images of bright field (BF), GFP, and RFP images, and lower panels are merged images of GFP and RFP images. The dashed line in the first image of the lower panels denotes the outline of the pollen tube, and the time indicates the period elapsed post-bombardment (hours:minutes). Arrows and arrowheads indicate the vegetative and generative cell nuclei, respectively. See also Video S2. **b, c:** Pollen tube germination on pistils using pollen hydrated on agarose germination medium. After spreading on the agarose medium, pollen grains were collected and used to pollinate the emasculated stigmas of *N. benthamiana* (**b**) and *N. tabacum* (**c**) pistils, respectively. Pollinated pistils were collected 24 h after pollination and stained with aniline blue solution. Bright field (BF) and ultraviolet (UV) illuminated images were merged. **d:** Semi-*in vivo* analysis of pollen tubes in *N. tabacum* pollinated after bombardment. Among the pollen tubes that emerged from the cut end of a pollinated pistil, two fluorescent-positive pollen tubes were observed. Arrows indicate fluorescent signals in the vegetative nuclei. Magnified images are also shown. VN, vegetative nucleus; GCN, generative cell nucleus. Scale bars: **a:** 20 μm, **b, c:** 500 μm, **d:** 100 μm.

**Fig. 5.**
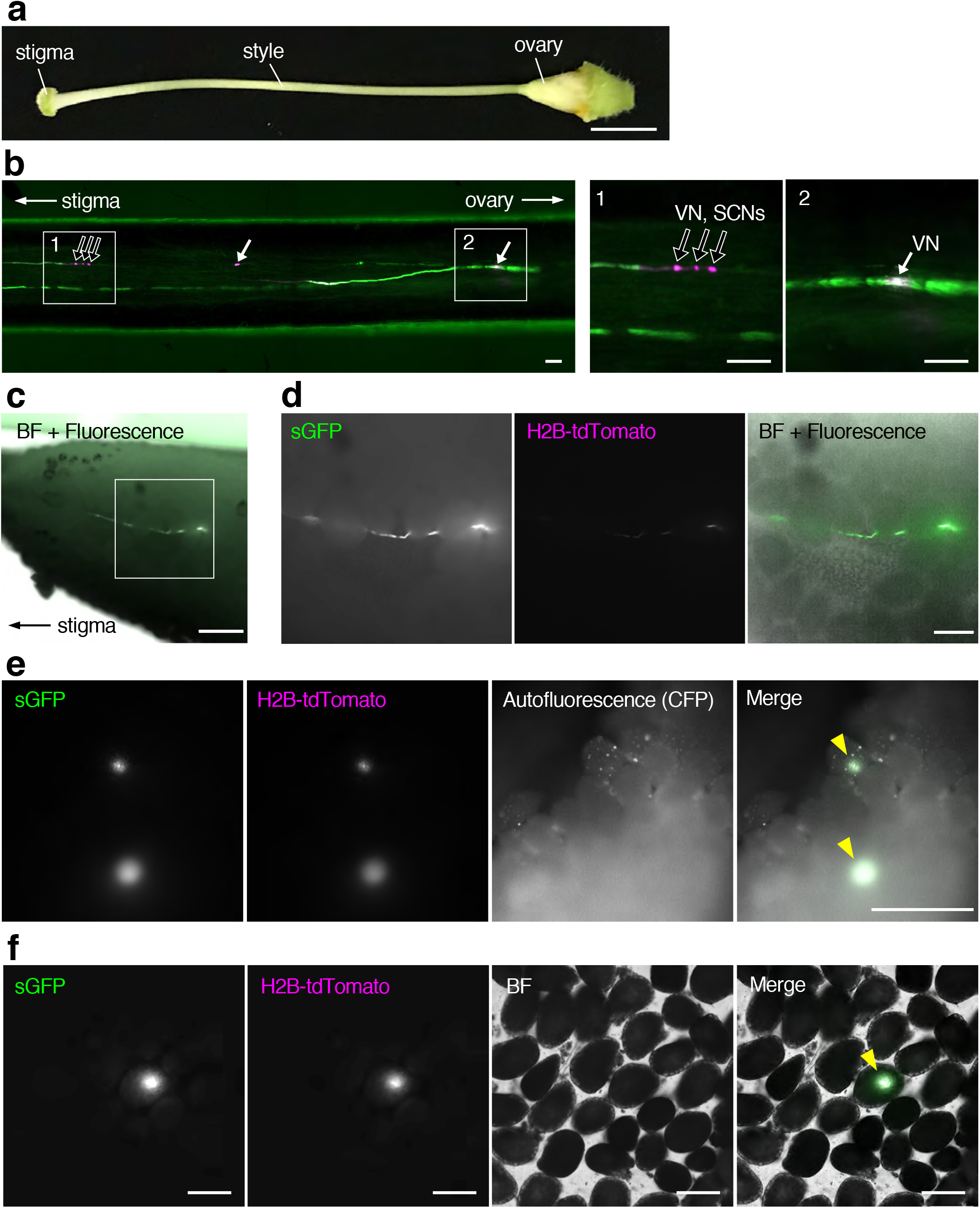
*In vivo* pollination of bombarded pollen with *AtUBQ10p::sGFP* and *35Sp::H2B-tdTomato* plasmid DNAs in *N. tabacum*. **a:** The pistil of *N. tabacum*. **b:** Vertical section of a style with bombarded pollen at 24 h after pollination. Pollinated style was hand dissected by tweezers and placed on the 10% glycerol. Magnified images were shown in the right panel. **c:** Pollen tubes from the bombarded pollen on the placenta 48 h after pollination. Ovary wall was removed by tweezers and placed on the 10% glycerol. Magnified images are shown in **d**. **e:** Two ovules on the placenta that received pollen tube from the bombarded pollen 48 h after pollination. Ovary wall was removed by tweezers and placed on 10% glycerol. **f:** Dissected ovules on 10% glycerol from the placenta at 48 h after pollination with bombarded pollen. Single ovule that receives pollen tube from the bombarded pollen. White arrows indicate the vegetative nucleus; black arrows show vegetative and generative nuclei. Yellow arrowheads indicate the ovule showing fluorescent signal derived from the bombarded pollen. VN, vegetative nucleus; GCN, generative cell nucleus; SCN, sperm cell nucleus. Scale bars: **a, c, e:** 500 μm, **b, f:** 100 μm, **d:** 200 μm.

## Discussion

Animal germ cells, such as those of sperm, eggs, zygotes, and embryos, contain a genome that develops into the individuals of the next generation. To obtain gene-modified organisms by inducing heritable genetic changes, genome editing has been studied using germline cells in animals (A. Lea and K. Niakan 2019; Cooper et al. 2017; Koslova et al. 2020). With respect to plants, somatic cells are generally used for genome engineering since that retain totipotency (Ikeuchi et al. 2015). Recently, plant germ cells, such as the zygotes of rice (Toda et al. 2019), have also been used to produce genome-modified plants. However, sophisticated anatomical techniques are required to isolate zygotes, as female germ cells are embedded deep within tissues (Mizuta and Higashiyama 2018). Moreover, tissue culture is necessary to generate genome-edited plants subsequent to zygote delivery. Compared with female germ cells, pollen is readily isolated and has a simple structure comprising two sperm cells. Pollen can thus be used as a “vector” for gene engineering of plant germ cells via fertilization (Resch and Touraev 2010). With regard to the number of cells contained within pollen grains, angiosperm pollen is generally divided into two types: bi-cellular and tri-cellular (Russell and Jones 2015). Approximately 30% of angiosperms produce tri-cellular pollen comprising two sperm cells at anthesis (Brewbaker 1967). The other 70% of angiosperms, including *N. benthamiana* and *N. tabacum*, produce bi-cellular pollen grains that contain a generative cell as a precursor of sperm cells. After anthesis, the generative cell undergoes mitosis to form two sperm cells within the pollen tube (Hackenberg and Twell 2019). These two sperm cells fertilize an egg cell and a central cell, respectively (Berger et al. 2008). Consequently, only genetic material in the sperm cell that fertilizes the egg cell is transmitted to the progeny. In the present study, we detected the expression of delivered fluorescent proteins 2 to 3 h post-bombardment (Fig. 3a and Video S1) and division into two sperm cells was observed at approximately 10 h (Fig. 2d)—results that are comparable to observations of pollen tubes derived from non-bombarded wild-type pollen (Tian et al. 2005). Expression of the Cas9 protein is typically detectable within 5 h after transfection (Fajrial et al. 2020), and the findings of the present study indicate that when generative cells are mutated by biolistic delivery of CRISPR/Cas9, it is possible that the mutation is inherited in both sperm cells in bi-cellular pollen. Bi-cellular pollen is accordingly considered to be more suitable than tri-cellular pollen for genome engineering of sperm cells in pollen tubes. In general, the DNA repair pathway is closely associated with the cell cycle, in which it plays a key role in maintaining genomic integrity during mitosis (Mao et al. 2008). The cell cycle of pollen vegetative cells enters the G_0_ phase, whereas at anthesis the cell cycle of the generative cell is in the G_1_ S, or G_2_ phase, depending on the species (Borg and Twell 2010; Friedman 1999). Our observations in the present study indicate that the efficiency of genome editing in the pollen tube is lower than that in leaves, which could be attributed to the fact that the cells undergo only a single round of mitosis or that the cell cycle of the generative cell is at the late interphase. Moreover, this may be due to chromatin condensation of the generative cell.

The initial step in plant genetic engineering is the delivery of genes into plant cells. However, the protective rigid cell walls of most angiosperms tend to limit the effective delivery of most molecular types. Conventional methods used to deliver genes into plant cells can be grouped into three categories, namely physical, chemical, or biological (Birch 1997; Han and Kim 2019). Among these, one of the most commonly used biological approaches for generating gene-modified plants is *Agrobacterium*-mediated transformation (Bevan 1984). However, for many species, including those of economic importance, a drawback of this technique is the prerequisite of tissue culturing steps to facilitate plant regeneration. As an alternative, a physical gene delivery approach, biolistic particle delivery (also referred to as particle bombardment or gene gun delivery), has been widely used in plant species, including non-model plants (Sachin Rustgi 2020). This method, which is effective regardless of species or tissue type, entails a simple rapid procedure that can efficiently deliver a range of molecular types, including DNA, RNA, proteins, and dyes (Martin-Ortigosa and Wang 2014; Wang and Jiang 2011; Zhang et al. 2016). In this regard, when seeking to introduce DNA, such as plasmid vectors, a promoter that functions stably in the introduced cells is required. For example, whereas the *AtRPS5A* promoter has been shown to be efficient for driving the expression of Cas9 in *A. thaliana* germ cells via *Agrobacterium*-mediated transformation (Ordon et al. 2020; Tsutsui and Higashiyama 2017), we found that it tends to be unstable when delivered via particle bombardment in both *N. benthamiana* and *N. tabacum* pollen tubes (Table 1 and Figure 1a). In contrast, we observed that the *AtUBQ10* promoter showed high and stable activity in the pollen tube of the four species examined in the present study (Table 1). Under transient conditions, the CaMV 35S promoter showed activity in tobacco pollen, but not in either torenia or tomato pollen (Table 1). By using transformants, it is known that the CaMV 35S promoter is not active in *Arabidopsis* pollen, whereas it is active in tobacco pollen (Wilkinson et al. 1997). CaMV 35S promoter activity in pollen may differ among species. It is thus conceivable that the *AtUBQ10* promoter sequence contains a universal sequence that enhances the efficiency of genome editing in pollen tubes (Zheng et al. 2020).

Although biolistic delivery is typically used to facilitate transient expression, methods for producing transformants or genome-edited plants from bombarded cells have also been reported. Recent studies have reported successful editing of plant genetic material via particle bombardment delivery of plasmids, *in vitro* transcripts, or ribonucleoprotein complexes (RNPs) of CRISPR/Cas9 complexes using wheat and maize embryo-derived callus (Liang et al. 2018; Svitashev et al. 2016; Svitashev et al. 2015). Furthermore, for wheat, an *in planta* transformation method using biolistic delivery has been reported that does not require callus culture and regeneration (Hamada et al. 2017). In the case of pollen, transformants produced via regeneration from bombarded pollen have also been reported (Stöger et al. 1995). In tobacco, only five antibiotic-resistant seeds harboring transgenes were defined from 30,000 obtained seeds (Touraev et al. 1997). Although, in the present study, we succeeded in introducing genes into pollen via particle bombardment, the efficiency achieved was typically low. We detected a total of 11 ovules showing fluorescent expression from four ovaries pollinated with bombarded pollen (Fig. 5e, 5f). Given that the number of seeds in *N. tabacum* is approximately 530–1,000 per ovary (Touraev et al. 1997), it is estimated that the percentage of fluorescent expressed ovules is approximately 0.275−0.519% of the total. The delivery into the generative cell is approximately one sixth, and it is estimated that 0.045−0.087% of the total ovules are derived from bombarded pollen introduced into the generative cell. Genome editing occurs only in a fraction of the bombarded pollen, and only a few of these may subsequently achieve successful fertilization. Therefore, it is expected that the number of seeds containing edited genomes will be extremely small. Additionally, when genome editing occurs in pollen and the plasmid DNA is not delivered to the egg, the target locus in the resulting seed is expected to be heterozygous. Therefore, it is essential to be able to efficiently detect or select genetically modified seeds among a large excess of unmodified seeds. On the other hand, continuous efforts are critical for the efficient production of genetically modified seeds by biolistic delivery. Efficient production of genome-edited plants will require both efficient biolistic delivery and improved efficacy of genome editing. With regard to the former, our detection method of biolistic delivery in pollen and pollinated pistils would contribute to increasing the efficiency at various reproductive steps. In the latter regard, a number of approaches aimed at enhancing the efficiency of genome editing have been reported, including the modification of Cas9 (Ling et al. 2020; Osakabe et al. 2020), design of guide RNA (Moon et al. 2019), the use of RNPs (Liang et al. 2018; Svitashev et al. 2016), and the use of small chemical compounds (Yu et al. 2015). Various selection methods have also been reported, including those based on antibiotic resistance (Chesnokov and Manteuffel 2000) and herbicide resistance, by targeting endogenous genes (Han and Kim 2019). These applications and our detection method will contribute to further advances in the engineering of plant genomes, including those of economically important crops and non-model plants.

## Declarations

## Availability of data and material

The authors confirm that the data supporting the findings of this study are available within the article and its supplementary materials.

## Code availability

Not applicable

## Ethics approval

Not applicable

## Consent to participate

Approved

## Consent for publication

Approved

## Key Message

Biolistic delivery into pollen

## Short legends for supporting information

**Table S1** List of the plasmid DNA vectors used in this study

**Table S2** List of the primers used in this study

**Video S1** Time-lapse images of fluorescent protein expression in the bombarded pollen. See also Figure 3a. The times denote the period elapsed post-bombardment (hours:minutes). Arrows and arrowheads indicate mClover in the nucleus and the tip of pollen tube of the bombarded pollen.

**Video S2** Movement of the generative cell nucleus in a pollen tube germinated from bombarded pollen. See also Figure 4a. The times denote the period elapsed post-bombardment (hours:minutes). Arrows and arrowheads indicate the vegetative and generative cell nuclei, respectively.

## Acknowledgements

We sincerely thank D. Kurihara, M. Ueda, and N. Inada for providing vectors; M. Notaguchi and the Leaf Tobacco Research Center, Japan Tobacco, Inc. for providing seeds; H. Adachi and H. Yoshioka for advising transformation; M. Fujii, T. Shinagawa, M. Igarashi, and Y. Hiramatsu for preparing plant materials and performing experiments. This work was supported by the Japan Science and Technology Agency (JST) [Precursory Research for Embryonic Science and Technology (PRESTO); grant number JPMJPR15QC] to Y.M. and the Japan Society for the Promotion of Science [Grant-in-Aid for Transformative Research Areas (KAKENHI); grant No. 20H05778, 20H05779] to Y.M.

## References

A. Lea R, K. Niakan K (2019) Human germline genome editing. Nature Cell Biology 21:1479–1489 doi:10.1038/s41556-019-0424-0

Adachi S et al. (2011) Programmed induction of endoreduplication by DNA double-strand breaks in Arabidopsis. Proceedings of the National Academy of Sciences 108:10004–10009 doi:10.1073/pnas.1103584108

Berger F, Hamamura Y, Ingouff M, Higashiyama T (2008) Double fertilization - caught in the act. Trends Plant Sci 13:437–443 doi:doi.org/10.1016/j.tplants.2008.05.011

Bevan M (1984) Binary Agrobacterium vectors for plant transformation. Nucleic Acids Research 12:8711–8721 doi:10.1093/nar/12.22.8711

Bhowmik P et al. (2018) Targeted mutagenesis in wheat microspores using CRISPR/Cas9. Scientific reports 8:6502 doi:10.1038/s41598-018-24690-8

Birch RG (1997) PLANT TRANSFORMATION: Problems and Strategies for Practical Application. Annual Review of Plant Physiology and Plant Molecular Biology 48:297–326 doi:10.1146/annurev.arplant.48.1.297

Borg M, Twell D (2010) Life after meiosis: patterning the angiosperm male gametophyte. Biochemical Society transactions 38 2:577–582 doi:doi.org/10.1042/BST0380577

Bregitzer P, Halbert SE, Lemaux PG (1998) Somaclonal variation in the progeny of transgenic barley. Theoretical and Applied Genetics 96:421–425 doi:10.1007/s001220050758

Brewbaker JR (1967) The distribution and phylogenetic significance of binucleate and trinucleate pollen grains in the angiosperms. American journal of botany 54 doi:10.2307/2440530

Chesnokov YV, Manteuffel R (2000) Kanamycin resistance of germinating pollen of transgenic plants. Sexual Plant Reproduction 12:232–236 doi:10.1007/s004970050006

Clough SJ, Bent AF (1998) Floral dip: a simplified method for Agrobacterium -mediated transformation of Arabidopsis thaliana. The Plant Journal 16:735–743 doi:10.1046/j.1365-313x.1998.00343.x

Cong L et al. (2013) Multiplex genome engineering using CRISPR/Cas systems. Science 339:819–823 doi:doi:10.1126/science.1231143

Cooper CA et al. (2017) Generation of gene edited birds in one generation using sperm transfection assisted gene editing (STAGE). Transgenic research 26:331–347 doi:10.1007/s11248-016-0003-0

Dresselhaus T, Sprunck S, Wessel GM (2016) Fertilization Mechanisms in Flowering Plants. Current biology: CB 26:R125–139 doi:doi.org/10.1016/j.cub.2015.12.032

Eady C, Lindsey K, Twell D (1995) The Significance of Microspore Division and Division Symmetry for Vegetative Cell-Specific Transcription and Generative Cell Differentiation. The Plant cell 7:65–74 doi:10.1105/tpc.7.1.65

Eapen S (2011) Pollen grains as a target for introduction of foreign genes into plants: an assessment. Physiology and molecular biology of plants: an international journal of functional plant biology 17:1–8 doi:10.1007/s12298-010-0042-6

Fajrial AK, He QQ, Wirusanti NI, Slansky JE, Ding X (2020) A review of emerging physical transfection methods for CRISPR/Cas9-mediated gene editing. Theranostics 10:5532–5549 doi:10.7150/thno.43465

Fossi M, Amundson K, Kuppu S, Britt A, Comai L (2019) Regeneration of Solanum tuberosum Plants from Protoplasts Induces Widespread Genome Instability. Plant physiology 180:78–86 doi:10.1104/pp.18.00906

Friedman WE (1999) Expression of the cell cycle in sperm of Arabidopsis: implications for understanding patterns of gametogenesis and fertilization in plants and other eukaryotes. Development (Cambridge, England) 126:1065–1075

Hackenberg D, Twell D (2019) Chapter Eleven - The evolution and patterning of male gametophyte development. In: Grossniklaus U (ed) Current Topics in Developmental Biology, vol 131. Academic Press, pp 257–298. doi:10.1016/bs.ctdb.2018.10.008

Hamada H, Linghu Q, Nagira Y, Miki R, Taoka N, Imai R (2017) An in planta biolistic method for stable wheat transformation. Scientific reports 7:11443 doi:10.1038/s41598-017-11936-0

Han Y-J, Kim J-I (2019) Application of CRISPR/Cas9-mediated gene editing for the development of herbicide-resistant plants. Plant Biotechnology Reports 13:447–457 doi:10.1007/s11816-019-00575-8

Hoekstra FA, Bruinsma J (1975) Respiration and Vitality of Binucleate and Trinucleate Pollen. Physiologia Plantarum 34:221–225 doi:10.1111/j.1399-3054.1975.tb03825.x

Ikeuchi M et al. (2015) PRC2 represses dedifferentiation of mature somatic cells in Arabidopsis. Nature Plants 1:15089 doi:10.1038/nplants.2015.89

Koslova A et al. (2020) Precise CRISPR/Cas9 editing of the NHE1 gene renders chickens resistant to the J subgroup of avian leukosis virus. Proceedings of the National Academy of Sciences of the United States of America 117:2108–2112 doi:10.1073/pnas.1913827117

Li J-F et al. (2013) Multiplex and homologous recombination-mediated genome editing in Arabidopsis and Nicotiana benthamiana using guide RNA and Cas9. Nat Biotech 31:688–691 doi:10.1038/nbt.2654

Liang Z, Chen K, Zhang Y, Liu J, Yin K, Qiu JL, Gao C (2018) Genome editing of bread wheat using biolistic delivery of CRISPR/Cas9 in vitro transcripts or ribonucleoproteins. Nature protocols 13:413–430 doi:10.1038/nprot.2017.145

Ling X et al. (2020) Improving the efficiency of precise genome editing with site-specific Cas9-oligonucleotide conjugates. Science Advances 6:eaaz0051 doi:10.1126/sciadv.aaz0051

Mao Z, Bozzella M, Seluanov A, Gorbunova V (2008) DNA repair by nonhomologous end joining and homologous recombination during cell cycle in human cells. Cell Cycle 7:2902–2906 doi:10.4161/cc.7.18.6679

Martin-Ortigosa S, Wang K (2014) Proteolistics: a biolistic method for intracellular delivery of proteins. Transgenic research 23:743–756 doi:10.1007/s11248-014-9807-y

Mizuta Y, Higashiyama T (2018) Chemical signaling for pollen tube guidance at a glance. J Cell Sci 131 doi:10.1242/jcs.208447

Mizuta Y, Kurihara D, Higashiyama T (2015) Two-photon imaging with longer wavelength excitation in intact *Arabidopsis* tissues. Protoplasma 252:1231–1240 doi:10.1007/s00709-014-0754-5

Moon SB, Kim DY, Ko J-H, Kim J-S, Kim Y-S (2019) Improving CRISPR Genome Editing by Engineering Guide RNAs. Trends in biotechnology 37:870–881 doi:10.1016/j.tibtech.2019.01.009

Nekrasov V, Staskawicz B, Weigel D, Jones JDG, Kamoun S (2013) Targeted mutagenesis in the model plant Nicotiana benthamiana using Cas9 RNA-guided endonuclease. Nat Biotech 31:691–693 doi:10.1038/nbt.2655

Newell CA (2000) Plant transformation technology. Molecular Biotechnology 16:53–65 doi:10.1385/mb:16:1:53

Okuda S et al. (2009) Defensin-like polypeptide LUREs are pollen tube attractants secreted from synergid cells. Nature 458:357–361 doi:10.1038/nature07882

Ordon J, Bressan M, Kretschmer C, Dall’Osto L, Marillonnet S, Bassi R, Stuttmann J (2020) Optimized Cas9 expression systems for highly efficient Arabidopsis genome editing facilitate isolation of complex alleles in a single generation. Funct Integr Genomics 20:151–162 doi:10.1007/s10142-019-00665-4

Osakabe K et al. (2020) Genome editing in plants using CRISPR type I-D nuclease. Communications Biology 3:648 doi:10.1038/s42003-020-01366-6

Osakabe Y, Osakabe K (2015) Genome editing with engineered nucleases in plants. Plant & cell physiology 56:389–400 doi:10.1093/pcp/pcu170

Osakabe Y, Watanabe T, Sugano SS, Ueta R, Ishihara R, Shinozaki K, Osakabe K (2016) Optimization of CRISPR/Cas9 genome editing to modify abiotic stress responses in plants. Scientific reports 6:26685 doi:10.1038/srep26685

Palanivelu R, Preuss D (2006) Distinct short-range ovule signals attract or repel *Arabidopsis thaliana* pollen tubes *in vitro*. BMC Plant Biol 6:7 doi:10.1186/1471-2229-6-7

Paungfoo-Lonhienne C, Lonhienne TG, Mudge SR, Schenk PM, Christie M, Carroll BJ, Schmidt S (2010) DNA is taken up by root hairs and pollen, and stimulates root and pollen tube growth. Plant physiology 153:799–805 doi:10.1104/pp.110.154963

Resch T, Touraev A (2010) Pollen Transformation Technologies. In: Plant Transformation Technologies. pp 83–91. doi:10.1002/9780470958988.ch5

Russell SD, Jones DS (2015) The male germline of angiosperms: repertoire of an inconspicuous but important cell lineage. Frontiers in plant science 6:173–173 doi:10.3389/fpls.2015.00173

Sachin Rustgi HL (2020) Biolistic DNA Delivery in Plants. Methods in Molecular Biology. Humana Press. doi:1071603558

Sanford JC (2000) The development of the biolistic process. In Vitro Cellular & Developmental Biology - Plant 36:303–308 doi:10.1007/s11627-000-0056-9

Stöger E, Fink C, Pfosser M, Heberle-Bors E (1995) Plant transformation by particle bombardment of embryogenic pollen. Plant Cell Reports 14:273–278 doi:10.1007/BF00232027

Svitashev S, Schwartz C, Lenderts B, Young JK, Mark Cigan A (2016) Genome editing in maize directed by CRISPR–Cas9 ribonucleoprotein complexes. 7:13274 doi:10.1038/ncomms13274

Svitashev S, Young JK, Schwartz C, Gao H, Falco SC, Cigan AM (2015) Targeted Mutagenesis, Precise Gene Editing, and Site-Specific Gene Insertion in Maize Using Cas9 and Guide RNA. Plant physiology 169:931–945 doi:10.1104/pp.15.00793

Tian HQ, Yuan T, Russell SD (2005) Relationship between double fertilization and the cell cycle in male and female gametes of tobacco. Sexual Plant Reproduction 17:243–252 doi:10.1007/s00497-004-0233-9

Toda E et al. (2019) An efficient DNA- and selectable-marker-free genome-editing system using zygotes in rice. Nature Plants 5:363–368 doi:10.1038/s41477-019-0386-z

Touraev A, Stöger E, Voronin V, Heberle-Bors E (1997) Plant male germ line transformation. The Plant Journal 12:949–956 doi:10.1046/j.1365-313X.1997.12040949.x

Tsutsui H, Higashiyama T (2017) pKAMA-ITACHI vectors for highly efficient CRISPR/Cas9-mediated gene knockout in Arabidopsis thaliana. Plant Cell Physiol 58:46–56 doi:10.1093/pcp/pcw191

Wang H, Jiang L (2011) Transient expression and analysis of fluorescent reporter proteins in plant pollen tubes. Nature protocols 6:419–426 doi:10.1038/nprot.2011.309

Wang H, Yang H, Shivalila CS, Dawlaty MM, Cheng AW, Zhang F, Jaenisch R (2013) One-Step Generation of Mice Carrying Mutations in Multiple Genes by CRISPR/Cas-Mediated Genome Engineering. Cell 153:910–918 doi:10.1016/j.cell.2013.04.025

Wilkinson JE, Twell D, Lindsey K (1997) Activities of CaMV 35S and nos promoters in pollen: implications for field release of transgenic plants. Journal of experimental botany 48:265–275 doi:10.1093/jxb/48.2.265

Woo JW et al. (2015) DNA-free genome editing in plants with preassembled CRISPR-Cas9 ribonucleoproteins. Nat Biotech 33:1162–1164 doi:10.1038/nbt.3389

Yu C et al. (2015) Small Molecules Enhance CRISPR Genome Editing in Pluripotent Stem Cells. Cell Stem Cell 16:142–147 doi:10.1016/j.stem.2015.01.003

Zhang Y et al. (2016) Efficient and transgene-free genome editing in wheat through transient expression of CRISPR/Cas9 DNA or RNA. Nature Communications 7:12617 doi:10.1038/ncomms12617

Zhang Y, Zhang B, Yang T, Zhang J, Liu B, Zhan X, Liang Y (2020) The GAMYB-like gene SlMYB33 mediates flowering and pollen development in tomato. Horticulture Research 7:133 doi:10.1038/s41438-020-00366-1

Zhao X et al. (2017) Pollen magnetofection for genetic modification with magnetic nanoparticles as gene carriers. Nature Plants doi:10.1038/s41477-017-0063-z

Zheng X et al. (2020) The Improvement of CRISPR-Cas9 System With Ubiquitin-Associated Domain Fusion for Efficient Plant Genome Editing. Frontiers in plant science 11 doi:10.3389/fpls.2020.00621

Zhou L-z, Dresselhaus T (2019) Chapter Seventeen - Friend or foe: Signaling mechanisms during double fertilization in flowering seed plants. In: Grossniklaus U (ed) Current Topics in Developmental Biology, vol 131. Academic Press, pp 453–496. doi:10.1016/bs.ctdb.2018.11.013

Zuberi MI, Dickinson HG (1985) Pollen-stigma interaction in Brassica. III. Hydration of the pollen grains. Journal of Cell Science 76:321–336

