## Supplementary material for "Detection of a biolistic delivery of fluorescent markers and CRISPR/Cas9 to the pollen tube": TableS1

Table S1. List of plasmid DNA vectors used in this study

| Plasmid name | Description | Purpose | Related figure | Reference |
|---|---|---|---|---|
| DKv327 | *35Sp::mTFP1* | Evaluation of promoter activity | Figure 1a | This plasmid was provided from Dr. Noriko Inada. |
| DKv744 | *35Sp::H2B-tdTomato* | Evaluation of promoter activity, visualization of bombarded pollen t | Figure 1a, 2c, 4a, 4d, 5b-5g | This plasmid was provided from Dr. Daisuke Kurihar |
| sSNv26 | *AtUBQ10p::tdTomato* | Evaluation of promoter activity, visualization of bombarded pollen t | Figure 1a, 3a, 3c | Constructed in this study |
| sSNv10 | *AtRPS5Ap::sGFP* | Evaluation of promoter activity | Figure 1a | Constructed in this study |
| DKv277 | *AtRPS5Ap::H2B-tdTomato* | Evaluation of promoter activity, visualization of bombarded pollen t | Figure 1a | Adachi et al, 2011 |
| sSNv28 | *AtUBQ10p::H2B-mClover* | Transformation, visualization of bombarded pollen tube | Figure 1a, 2a, 2b, 2d, 2f, 3a, 4a, 4d | Constructed in this study |
| YMv32 | *LAT52p::mApple* | Multiple expression, evaluation of delivery efficiency | Figure 2a, 2b , 4d | Mizuta et al., 2015 |
| sSNv21 | *AtUBQ10p::Cas9/U6.26p::NbPDS3-sgRNA* | Genome editing in leaf and pollen | Figure 3b, 3d | Constructed in this study |
| sSNv25 | *AtUBQ10p::sGFP* | *N. tabacum* semi-in vivo assay | Figure 4d, 5b-5g | Constructed in this study |
