## Supplementary material for "Detection of a biolistic delivery of fluorescent markers and CRISPR/Cas9 to the pollen tube": TableS2

Table S2. List of primers used in this study

| Primer name | 5' -> 3' sequence | Purpose |
|---|---|---|
| DKp113 | CCATGGTGAGCAAGGGCGAGGAG | Construction of sSNv10 |
| DKp130 | CCATGTCGACGGCTGTGGTGAGAG | Construction of sSNv10 |
| NbPDS3_sgRNA-1_FWD | ATTGCCGTTAATTTGAGAGTCCA | Construction of CRISPR/Cas9 vector targeting NbPDS3 |
| NbPDS3_sgRNA-1_RVS | AAACTGGACTCTCAAATTAACGG | Construction of CRISPR/Cas9 vector targeting NbPDS3 |
| RPS5Ap-R1 | AGATGTGATGAACGCCACAG | Construction of modified pKI1.1 vector |
| pTTK346_inverse_F | CCATGGACTATAAGGACCACGAC | Construction of modified pKI1.1 vector |
| GA_UBQ10p_FWD | GCGTTCATCACATCTCGACGAGTCAGTAATAAACG | Construction of modified pKI1.1 vector |
| GA_UBQ10p_RVS | CCTTATAGTCCATGGCTGTTAATCAGAAAAACTCAG | Construction of modified pKI1.1 vector |
| PDS_MlyIF | GCTTTGCTTGAGAAAAGCTCTC | PCR of NbPDS3 sequence |
| PDS_MlyIR | ACATAACAAATTCCTTTGCAAGC | PCR of NbPDS3 sequence |
| NbPDS3_nest_F | TTTTCCCGTTTAGGATCTTG | Nested PCR of NbPDS3 sequence |
| NbPDS3_nest_R | GCAAACATCTTGACTTTCAG | Nested PCR of NbPDS3 sequence |
| M13 forward | GTAAAACGACGGCCAGT | Colony PCR to detect mutation in NbPDS3 |
| M13 reverse | CAGGAAACAGCTATGAC | Colony PCR to detect mutation in NbPDS3 |
| NbPDS3 primer-m | GATAAGCTGAATTACCTTGGAC | Colony PCR to detect mutation in NbPDS3 |
